# Effects of excitotoxicity in the hypothalamus in transgenic mouse models of Huntington disease

**DOI:** 10.1101/2021.04.08.439100

**Authors:** Jo B. Henningsen, Rana Soylu-Kucharz, Maria Björkqvist, Åsa Petersén

## Abstract

Huntington disease (HD) is a fatal neurodegenerative movement disorder caused by an expanded CAG repeat in the huntingtin gene (HTT). The mutant huntingtin protein is ubiquitously expressed, but only certain brain regions are affected. The hypothalamus has emerged as an important area of pathology with selective loss of neurons expressing the neuropeptides orexin (hypocretin), oxytocin and vasopressin in human postmortem HD tissue. Hypothalamic changes in HD may have implications for early disease manifestations affecting the regulation of sleep, emotions and metabolism. The underlying mechanisms of selective vulnerability of certain neurons in HD are not fully understood, but excitotoxicity has been proposed to play a role. Further understanding of mechanisms rendering neurons sensitive to mutant huntingtin may reveal novel targets for therapeutic interventions. In the present study, we wanted to examine whether transgenic HD mice display altered sensitivity to excitotoxicity in the hypothalamus. We first assessed effects of hypothalamic injections of the excitotoxin quinolinic acid (QA) into wild-type (WT) mice. We show that neuronal populations expressing melanin-concentrating hormone (MCH) and cocaine and amphetamine-regulated transcript (CART) display a dose-dependent sensitivity to QA. In contrast, neuronal populations expressing orexin, oxytocin, vasopressin as well as tyrosine hydroxylase in the A13 area are resistant to QA-induced toxicity. We demonstrate that the R6/2 transgenic mouse model expressing a short fragment of mutant HTT displays hypothalamic neuropathology with discrete loss of the neuronal populations expressing orexin, MCH, CART, and orexin at 12 weeks of age. The BACHD mouse model expressing full-length mutant HTT does not display any hypothalamic neuropathology at 2 months of age. There was no effect of hypothalamic injections of QA on the neuronal populations expressing orexin, MCH, CART or oxytocin in neither HD mouse model. In conclusion, we find no support for a role of excitotoxicity in the loss of hypothalamic neuronal populations in HD.

## Introduction

The underlying mechanisms of selective vulnerability of certain brain regions in the neurodegenerative Huntington disease are not known. HD is a fatal disorder caused by an expanded CAG repeat in the HTT gene. The disease-causing mutant huntingtin protein is ubiquitously expressed, yet the striatal medium spiny neurons (MSNs) in the basal ganglia are the most affected. Striatal atrophy is associated with the typical movement disorder that, together with a genetic test, leads to the clinical diagnosis. There is no disease-modifying therapy available, and affected patients progress in their disease leading to premature death around 20 years after diagnosis. Understanding the early pathogenesis of the disease may lead to the identification of novel targets for therapeutic intervention. Excitotoxicity, i.e., excessive glutamate receptor stimulation, has been suggested to be involved in the selective vulnerability of striatal neurons in HD. Excitotoxic striatal lesions produced by injections of quinolinic acid (QA), an endogenous agonist of the N-methyl-D-aspartate (NMDA) glutamate receptor, in wild-type rodents and primates have been shown to recapitulate a similar loss of MSNs as in HD ^1-4^. Transgenic mouse models of HD display altered striatal sensitivity to excitotoxicity. Early studies showed that the transgenic R6/1 and the R6/2 mouse models of HD, which express a short fragment of mutant HTT, displayed resistance to intrastriatal injection of QA ^5-7^. Other studies in mice expressing full-length mutant HTT, such as the yeast artificial chromosome (YAC)128 and the bacterial artificial chromosome (BAC) transgenic mouse model of HD (BACHD), showed increased sensitivity to intrastriatal injections of QA ^8,9^. Later studies have proposed that there is a biphasic response to excitotoxicity in HD with younger HD mice displaying increased sensitivity and older HD mice developing resistance to the insult as compared to wild-type littermates ^10,11^. Hence, there is an interplay between mutant HTT and the response of striatal neurons to excitotoxicity that may contribute to the selective vulnerability in HD.

Emerging studies indicate that also the hypothalamus is affected in HD, which may have implications for early non-motor features such as psychiatric symptoms, sleep disturbances and metabolic dysfunction ^12^. In clinical postmortem tissue from HD cases, there is a selective vulnerability of the neuronal populations expressing the neuropeptides orexin (hypocretin), oxytocin, and vasopressin ^13-17^. Hypothalamic pathology has also been recapitulated in the R6/2 and the YAC128 mouse models ^13,18-20^. Causal links between expression of mutant HTT in the hypothalamus and development of metabolic dysfunction as well as depressive-like behavior have been shown in the transgenic BACHD mice ^21,22^. The hypothalamus, including the lateral hypothalamic area (LHA) is, similarly to the striatum, rich in glutamatergic input from many other brain regions and is particularly enriched in NMDA receptors ^23^. NMDAR-NR2B subunit mRNA expression is increased in the hypothalamus of the R6/1 HD mouse model ^24^, implying that the glutamate homeostasis is altered in the hypothalamus in these mice. We hypothesized that excitotoxicity would play a role in the selective vulnerability of hypothalamic neurons in HD and therefore designed the present study to examine whether hypothalamic neurons, known to be affected in HD, would display altered sensitivity to excitotoxicity in transgenic mouse models of HD.

## Materials and Methods

### Animals

The experimental procedures were performed according to the approved guidelines by the local Animal Ethics Committee Malmö/Lund Animal Welfare and Ethics Committee (permits M65-13, M124-15 and 15499/2017). The mice were group-housed (2–5 animals) in universal Innocage mouse cages (InnoVive, San Diego, CA, USA) and kept under a 12 h light/dark cycle at 22 °C, with free access to water and standard chow diet. In this study, we carried out three sets of experiments. In the first part, we assessed effects of hypothalamic injections of the NMDA receptor agonist QA on the vulnerability of hypothalamic neuronal populations in WT mice from the FVB/NRj (hereafter referred to as FVB/N) strain (Janvier Labs). In the second part, we investigated effects of hypothalamic injections of QA in the transgenic R6/2 mouse model of HD. R6/2 and WT mice were obtained through crossing heterozygous males with WT females ^5^. Genotyping of mice was performed as described previously and sanger sequencing showed that the R6/2 mice used in this study had a 242-257 CAG repeat size, which results in a slower disease progression compared to R6/2 mice with 150 CAGs ^5^. In the third part of the study, we assessed effects of hypothalamic injections of QA in the BACHD mouse model of HD (FVB/N strain), which expresses full-length mutant HTT with 97 polyglutamine repeats ^25^. BACHD mice were obtained from the Jackson Laboratories (Bar Harbor, Maine, USA), and the colony was maintained by crossing male BACHD mice with WT FVB/N female mice. BACHD mice were genotyped from tail-tip samples using a conventional PCR procedure using primers for the BACHD transgene (forward primer: 5’-CCGCTCAGGTTCTGCTTTTA-3’ and reverse primer: 5’-AGGTCGGTGCAGAGGCTCCTC-3’) ^25^.

### Hypothalamic injections of quinolinic acid

On the day of injection, QA was dissolved in a 0.01 M PBS (pH 7.4) solution to the final concentration (ranging from 2.5 mM to 240 mM solution) and was kept on ice, while protected from light until use. The mice were anaesthetized using 2% isoflurane in oxygen/nitrous oxide (3:7) and received a subcutaneous analgesic injection (Temgesic, 0.1 mg/kg) prior to the surgical procedure. After adjusting for flat skull, the stereotactic coordinates used for injections into the dorsal lateral hypothalamic area were; 5 mm inferior to the dura matter, 0.5 mm anterior, and 0.8 mm lateral to Bregma. Using a pulled glass capillary (outer diameter app. 80 µm) adhered to a 5 µl Hamilton syringe (Reno, NV, USA), mice were unilaterally injected with either 0.5 µl of the QA solution or vehicle (saline), administered at a rate of 0.05 µl/15 s. Noteworthy, the stereotactic coordinates were optimized to prevent any physical damage of lateral hypothalamic cell populations. The capillary was left at the injection site for a minimum of 5 min post-injection to allow proper absorption of the QA and saline solutions and to prevent flow back upon capillary retraction.

### Immunohistochemistry

Seven days post-injection, mice were anesthetized with the terminal dose sodium pentobarbital and then transcardially perfused first with saline and then with pre-cooled 4% paraformaldehyde (PFA) at a 10 ml/min rate for 8 minutes. For post-fixation, brains were placed in 4% PFA solution at 4°C for 24 hours, after which PFA was replaced with a cold 25% sucrose solution for ∼24 hours. The brains were cut in six series of 30 μm thick coronal sections using a Microm HM450 microtome (Thermo Scientific). The sections were stored in an antifreeze solution at −20 ° (30% glycerol, 30% ethylene glycol solution in PBS). For immunohistochemistry, free-floating brain sections were rinsed 3 times for 10 minutes with 0.05 M Tris-buffered saline (TBS). Following that, a quenching reaction was performed in TBS containing 3% H_2_O_2_ and 10% Methanol. Brain sections were rinsed 3 times for 10 minutes in 0.25% triton-X in TBS (TBS-T). Next, the sections were pre-incubated with a blocking solution containing 5% normal goat serum and TBS-T for 1 hour at room temperature (RT). The sections were then incubated in the primary antibody solutions in TBS-T containing 5% normal goat serum at RT. The primary antibodies dilutions were as follows: anti-tyrosine hydroxylase (TH) (1:5000; anti-rabbit; Pel-Freez Biologicals; cat nb. P40101-50), anti-orexin (1:4000; anti-rabbit; Phoenix Pharmaceuticals; cat nb. H-003-30), anti-MCH (1:20000; anti-rabbit, Phoenix Pharmaceuticals; cat nb. H-070-47), anti-oxytocin (1:4000; anti-rabbit; Phoenix Pharmaceuticals; cat nb. H-051-01), anti-vasopressin (1:10000; anti-rabbit, MilliPore; cat nb. AB1565), anti-CART (1:4000; anti-rabbit; kindly provided by Dr. Kuhar). The next day, brain sections were rinsed 3 times for 10 minutes in TBS-T and then incubated for 1 hour in 1% bovine serum albumin TBS-T containing secondary biotinylated antibodies (1: 200; Vector Laboratories Inc.). Following the secondary antibody incubation step, the sections were washed 3 times for 10 minutes and staining was visualized using a 3,3’-diaminobenzidine (DAB) staining kit according to the manufacturer’s instructions (Vector Labs). Lastly, the chromatin-gelatin coated glass slides were used to mount the sections, and the slides were dried at RT. The sections were first dehydrated for the coverslipping in increasing alcohol solutions (70%, 95% and 100%) and then cleared in xylene. A Depex mounting medium (Sigma-Aldrich) was used for securing the sections.

### Stereological Analysis

The number of neuropeptidergic immunoreactive cells in the hypothalamus was assessed in a Nikon eclipse 80i light microscope using the VIS software (Visiopharm A/S) NewCast module, allowing the application of an unbiased counting frame in a design-based stereological approach. Quantifications were done using a 60x Plan-Apo oil objective, and all slides were blinded throughout the process. While all neuropeptide populations of interest were counted bilaterally, each hemisphere was analyzed separately according to the unilateral injection paradigm, allowing for comparison between injected versus un-injected hemispheres in vehicle as well as QA treated cohorts. The anatomical delineation of specific hypothalamic regions of interest was done in accordance with the Allen Mouse Brain Atlas. For quantification of orexin and MCH positive neurons in the LHA, the regions of interest (ROIs) were delineated to include all immunoreactive cells, corresponding to 6 sections/animal (orexin) and 7-8 sections/animal (MCH). ROIs for assessment of CART immunoreactive cells were limited to the lateral hypothalamic area, resulting in 6-7 sections/animal. Only TH positive cells within the Zona incerta (A13) were assessed in this study, corresponding to 4 sections/animal. Lastly, ROIs for detection of Vasopressin and Oxytocin positive cells were limited to the Paraventricular nucleus (PVN), resulting in 5 sections/animal (vasopressin) and 4-5 sections/animal (oxytocin). The fraction counted for each ROI was adjusted from 50%-100% to achieve a minimum of 200 cells counted/per brain. The quantitative output was adjusted for the number of sections (x6) and where adjusted for the % fraction counted, in order to estimate the total number of neurons per animal.

### Statistical Analysis

Statistical analyses were performed using Prism 8.0 (version 8.4.3, GraphPad). Data were tested first for normal distribution with the Shapiro-Wilk normality test. Next, the data were subjected to either the non-parametric Kruskal–Wallis followed by Dunn’s multiple-comparison test or Mann-Whitney test or the parametric one-way analysis of variance (ANOVA) followed by Tukey’s multiple-comparison test or Student’s t-test. The association between cell numbers (orexin and MCH) and increasing concentrations of QA was examined using simple linear regression analysis, and graphs were plotted with curves indicating a 95% confidence interval. For all data, the mean was provided with standard error of mean (SEM) in the results section. Data are plotted on the graphs as scatter dot plots, and bars represent mean ± SEM. In all tests, statistical differences were considered significant for p<0.05.

## Results and Discussion

### Effects of QA-induced excitotoxicity in the hypothalamus of wild-type mice

We began by determining the vulnerability to excitotoxicity in the hypothalamus in wild-type mice. For this purpose, we performed stereotaxic injections with increasing doses of the QA into the dorso-lateral part of the hypothalamus. Wild-type mice were injected unilaterally with increasing concentrations of QA (1.25 nmol – 120 nmol), including concentrations known to mediate substantial excitotoxic degeneration of striatal MSNs ^6^. Given that the hypothalamus is highly heterogeneous and that HD hypothalamic neuropathology is selective to certain hypothalamic populations ^12^, we assessed the effects on a selected number of neuronal populations in the injected as well as intact hemisphere seven days post injections. While the number of MCH-positive cells negatively correlated with increasing QA concentrations (Fig 1C-D), the number of orexin neurons was unaffected, despite a high QA concentration (120 nmol) (Fig 1A-B). The QA-mediated MCH neuron degeneration reached a plateau with QA concentrations ≥15 nmol, where we observed approximately 50% cell loss in the injected versus un-injected hemisphere (Fig 1C; F). Hence, the two major neuronal populations in the lateral hypothalamus, orexin and MCH, are not equally susceptible to QA-mediated excitotoxicity. This data is in agreement with a previous study performed in mice and rats where MCH neurons were sensitive to injections of QA whereas orexin neurons were not ^26^ and in contrast to an *in vitro* study on rat hypothalamic slices where QA exposure led to selective loss of orexin neurons ^27^.

**Figure 1.**
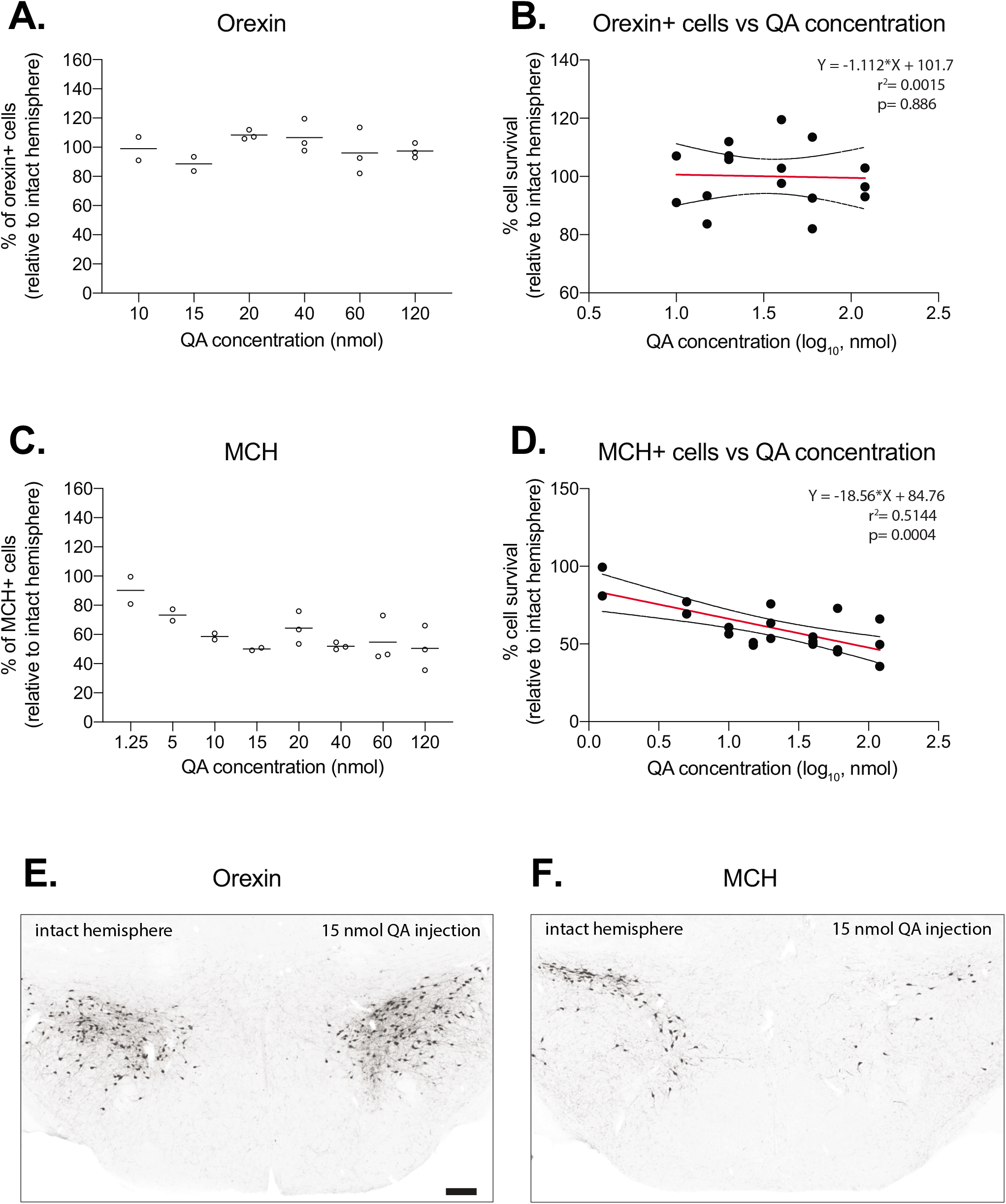
Effects of different concentrations of QA on orexin and MCH positive neurons in FVB/N wild-type mice. WT mice were injected with increasing concentrations of QA in the hypothalamus and examined seven days later. **(A)** Stereological estimation of orexin positive cells after exposure to intra-hypothalamic injections of increasing concentrations of QA. **(B)** There was no significant association between the percentage of orexin positive cells and increasing QA concentrations. **(C)** Assessment of MCH positive cells after exposure to increasing concentrations of QA. **(D)** There was a significant association between the percentage of MCH positive cells and increasing concentrations of QA. The data are presented as the number of orexin and MCH neurons in relation to the numbers in intact hemispheres. The dashed lines represent 95% confidence bands for the simple linear fit regression analysis, and QA concentrations are transformed to the base of Log10. Representative images of **(E)** orexin and **(F)** MCH immunohistochemistry showing the hypothalamus after unilateral injections of 15 nmol QA at 1-week post-injection. Scale bar = 200 μm.

We then continued to assess the effects of two concentrations of QA (15 nmol and 60 nmol) on four other neuropeptide expressing populations in the hypothalamus. The number of CART-positive neurons in the LHA was significantly reduced after a 60 nmol QA injection as compared to the intact hemispheres (Fig 2A and E, upper panel). In contrast, there was no effect of 15 nmol and 60 nmol QA on neither the TH positive A13 dopaminergic population of the zona incerta (Fig 2B) nor the oxytocin (Fig 2C) and AVP (Fig 2D) neurons located in the PVN.

**Figure 2.**
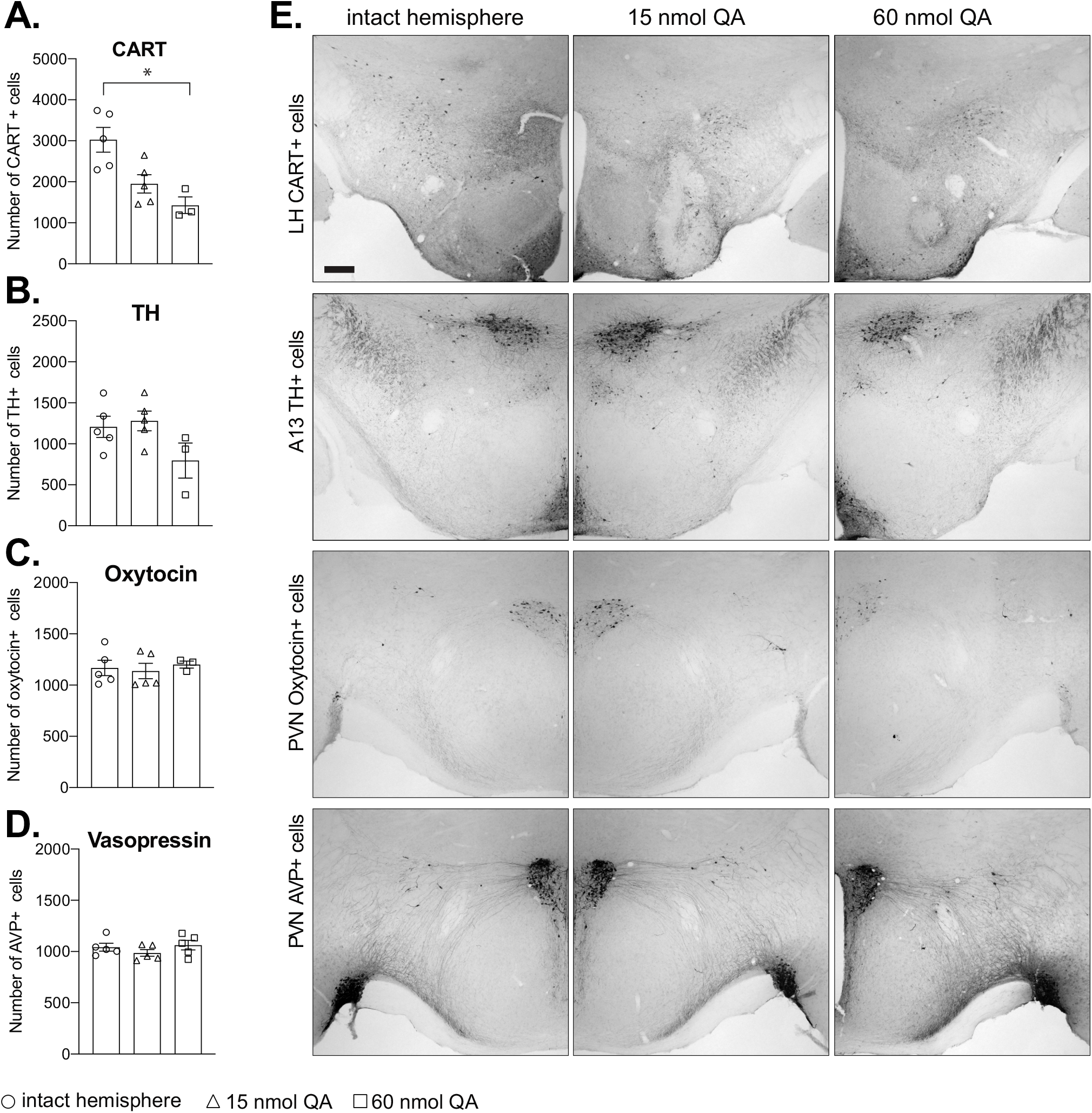
Stereological assessments of hypothalamic neuropeptide populations after exposure to QA. FVB/N WT mice were unilaterally injected with 15 nmol and 60 nmol of QA in the hypothalamus, and the number of neurons was examined seven days later on the injected side and compared to intact hemispheres of the 15 nmol QA group. **(A)** The number of CART-immunopositive cells was significantly reduced in the 60 nmol QA group compared to intact hemispheres (n=3-5/group, p=0.0136, Kruskal-Wallis test followed by Dunn’s multiple comparison test). There was no effect of injections of 15 nmol and 60 nmol QA on **(B)** A13 TH neurons, **(C)** oxytocin neurons in the paraventricular nucleus (PVN), and **(D)** vasopressin neurons in the PVN compared to the number in intact hemispheres (n=3-5/group, Kruskal-Wallis test followed by Dunn’s multiple comparison test). **(E)** Representative images are showing CART, TH, oxytocin, and vasopressin positive neurons in the hypothalamus. Data are represented as scatter dot plots, and bars represent mean ± SEM. Scale bar = 200 μm.

### Effects of QA-induced excitotoxicity in the hypothalamus of R6/2 HD mice

In the second part of the study, we aimed to investigate if hypothalamic vulnerability to excitotoxicity is altered in the R6/2 HD mouse model. We first determined the extent of hypothalamic neuropathology in the R6/2 ^(CAG 242-257)^ mice at an early (6 weeks) and advanced (12 weeks) symptomatic stage of the disease. In the present study, the R6/2 ^(CAG 242-257)^ mice display a significant loss of the number of orexin (Fig 3A), MCH (Fig 3B) and CART (Fig 3C) neurons localized in the lateral hypothalamic area, in addition to the loss of oxytocin neurons in the PVN (Fig 3D) at 12 weeks of age. The vasopressin-positive cell number is significantly reduced in R6/2 mice only at 6 weeks of age (Fig 3F), while the number of A13 TH-positive neurons is comparable between R6/2 ^(CAG 242-257)^ mice and controls at both 6 and 12 weeks (Fig 3E). This is in line with previous studies demonstrating loss of orexin ^13^, MCH ^19^, CART ^19,28^, oxytocin ^20,28^ and vasopressin ^20,28,29^ in the R6/2 mouse model.

**Figure 3.**
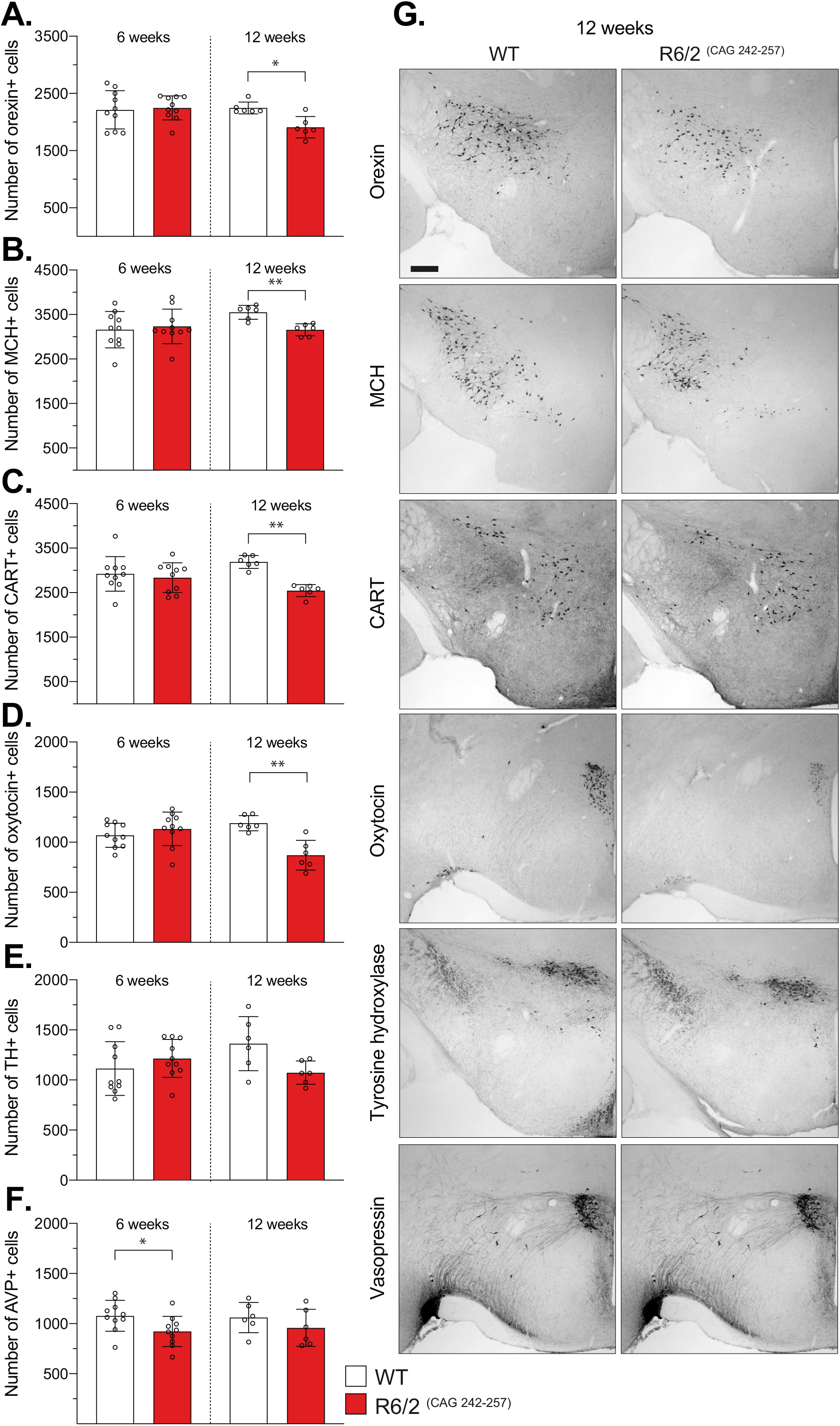
Stereological assessment of hypothalamic populations in the R6/2 mouse model expressing 242-257 CAG repeats in 6- and 12-weeks old mice. The number of **(A)** orexin (n=6/genotype, p=0.0260, Mann-Whitney test), **(B)** MCH (n=6/genotype, p=0.0022, Mann-Whitney test), **(C)** CART (n=6/genotype, p=0.0022, Mann-Whitney test) and **(D)** oxytocin (n=6/genotype, p=0.0043, Mann-Whitney test) positive cells were reduced at 12-week-old R6/2 mice compared to their WT littermate controls. **(E)** The number of A13 TH positive cells was comparable at both 6 and 12 weeks. **(F)** The vasopressin expressing cell numbers were significantly reduced in R6/2 mice compared to WT littermates at 6 weeks (n=10/genotype, two-tailed unpaired t-test, p=0.0341). **(F)** Representative photomicrographs of hypothalamic sections processed for orexin, MCH, CART, oxytocin, TH, and vasopressin immunohistochemistry in 12 weeks old WT and R6/2 mice. Data are represented as scatter dot plots, and bars represent mean ± SEM. Scale bar = 200 μm.

We then wanted to test whether R6/2 mice ^(CAG 242-257)^ displayed altered sensitivity to excitotoxicity compared to their wild-type littermates. Mice were therefore injected with 15 nmol QA into the hypothalamus at 6 and 12 weeks of age and assessed for hypothalamic neuropathology 7 days post-injection. At 6 weeks of age, there were no significant differences in the numbers of neurons expressing orexin, MCH, CART, oxytocin, vasopressin or TH independent of genotype or QA administration (Fig 4 A-F). At 12 weeks of age, there was only a significant reduction in MCH numbers in wild-type mice exposed to QA compared to controls (Fig 4 B) and no effect on the other examined neuronal populations of QA in wild-type mice. At this age, R6/2 mice exposed to QA displayed significantly different numbers of orexin, MCH, CART, and oxytocin immunopositive neurons compared to wild-type mice, but there were no significant differences to un-injected R6/2 mice (Fig 4). Hypothalamic injections of 15 nmol QA did not affect the numbers of vasopressin or TH-positive cells at any age or in any genotype. Taken together, hypothalamic injections of QA at a dose of 15 nmol do not exacerbate hypothalamic neuropathology in R6/2 mice at 6 or 12 weeks of age.

**Figure 4.**
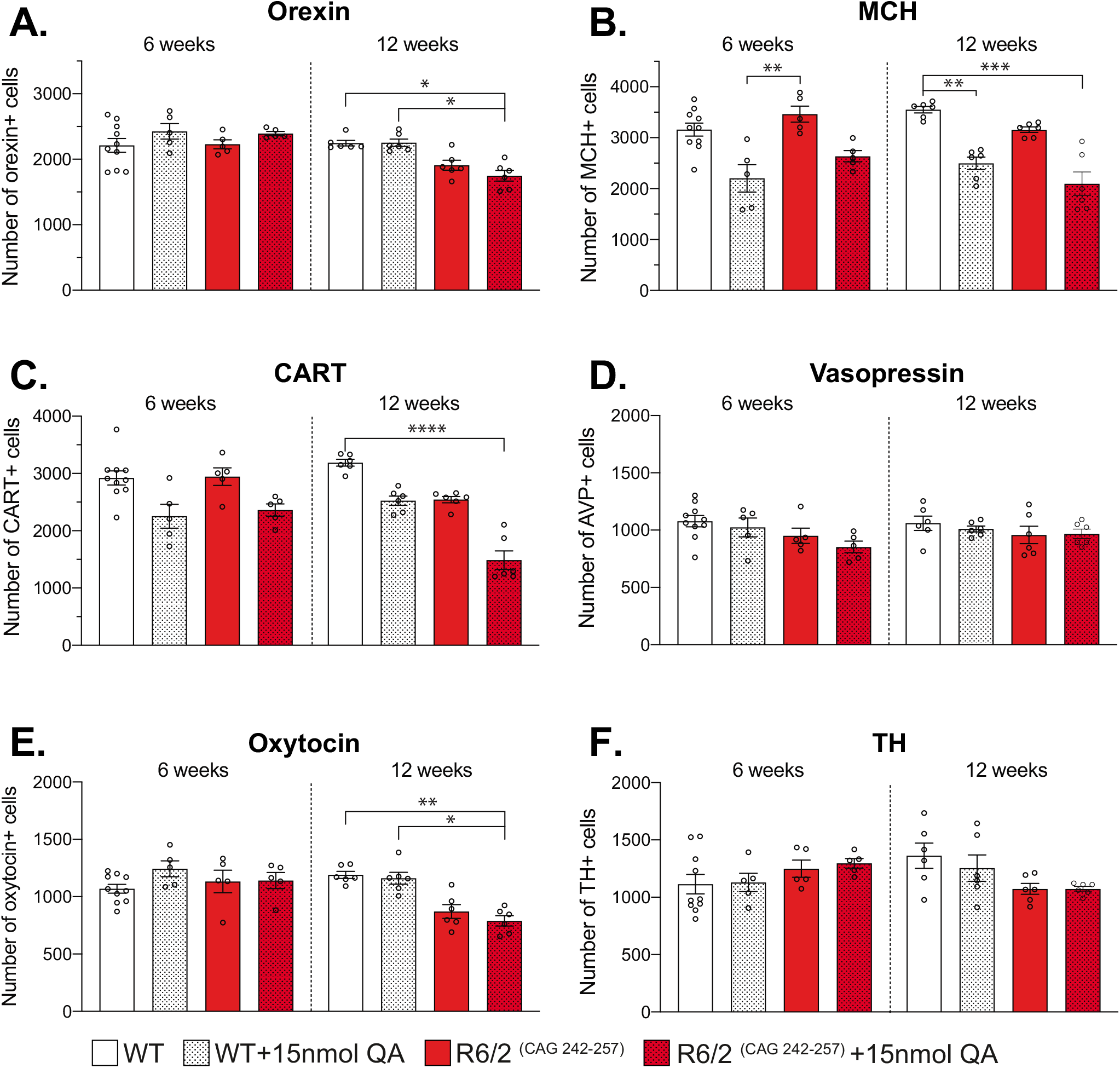
Assessment of neuronal populations in the R6/2 mice after hypothalamic injections of QA. The R6/2 mice and their WT littermate controls were unilaterally injected with 15 nmol QA in the hypothalamus at 6 and 12 weeks of age and hypothalamic neuropeptide expressing cells were examined after one week. **(A)** QA injections lead to a reduction of the orexin positive cell population in 12-week-old R6/2 mice compared to WT (p=0.0107) and WT-QA mice (n=6/group, p=0.0123). **(B)** The number of MCH positive cells was significantly reduced in WT-QA mice compared to the R6/2 mice at 6 weeks (n=5-10/group, p=0.0076). Both R6/2-QA (p=0.0004) and WT-QA (p=0.0049) mice had a significant reduction in the MCH cell population compared to WT mice at 12 weeks. **(C)** Stereological quantification of CART positive cells showed a decrease in 12-week-old R6/2 mice compared to WT mice (n=6/group, p<0.0001). **(D)** The hypothalamic injection of QA did not affect the vasopressin positive cell population at any age or genotype. **(E)** The number of oxytocin cells was significantly reduced in R6/2 mice injected with QA compared to WT (p=0.0161) and WT+QA group (n=6/group, p=0.0052) at 12 weeks. A Kruskal-Wallis test followed by Dunn’s multiple comparison statistical tests was performed. Data are represented as scatter dot plots, and bars represent mean ± SEM.

### No effect of QA-induced excitotoxicity in the hypothalamus of the BACHD mouse model of HD

In the final part of the study, we investigated whether the BACHD mouse model would display altered sensitivity to QA-induced excitotoxicity in the hypothalamus. Here, we included 2 months old BACHD mice to model the early and pre-symptomatic disease stage. Stereological assessment of the numbers of neuronal populations expressing orexin, MCH, CART or oxytocin showed no differences between 2 months old BACHD mice and wild-type littermate controls (Fig 5A). BACHD and wild-type control mice at 2 months of age were unilaterally injected with QA and examined 7 days post-injection. Based on reports of increased sensitivity to excitotoxicity in the striatum of pre-symptomatic HD models ^8,10^, a lower dose of QA (5 nmol) was included alongside two effective doses (15 nmol and 45 nmol) and the number of hypothalamic neuropeptide expressing cells were examined. When comparing wild-type and BACHD mice injected with QA, there was no effect of genotype on the number of orexin (Fig 5B), MCH (Fig 5C), CART (Fig 5D) or oxytocin (Fig 5E) immunoreactive cells with either of the QA concentrations investigated. Hence, BACHD mice at 2 months of age do not display altered sensitivity to excitotoxicity in the hypothalamus. However, considering the number of the animals used in the analysis (n=3-5), low statistical power could be a potential drawback of the study and resulting in false negatives. The response of hypothalamic neurons in the R6/2 and BACHD mouse models is different from the response reported to QA injections into the striatum of these models where striatal neurons show resistance or increased sensitivity to QA-mediated excitotoxicity ^7,9^. However, although several transgenic HD mouse models have displayed altered sensitivity to excitotoxicity in the striatum, there are also studies showing no signs of excitotoxicity in the striatum of a transgenic HD rat with around 22% of the huntingtin gene with 51 CAG repeats and another transgenic HD mouse model expressing 3 kb of mutant human huntingtin cDNA with 46 or 100 CAG repeats ^30,31^.

**Figure 5.**
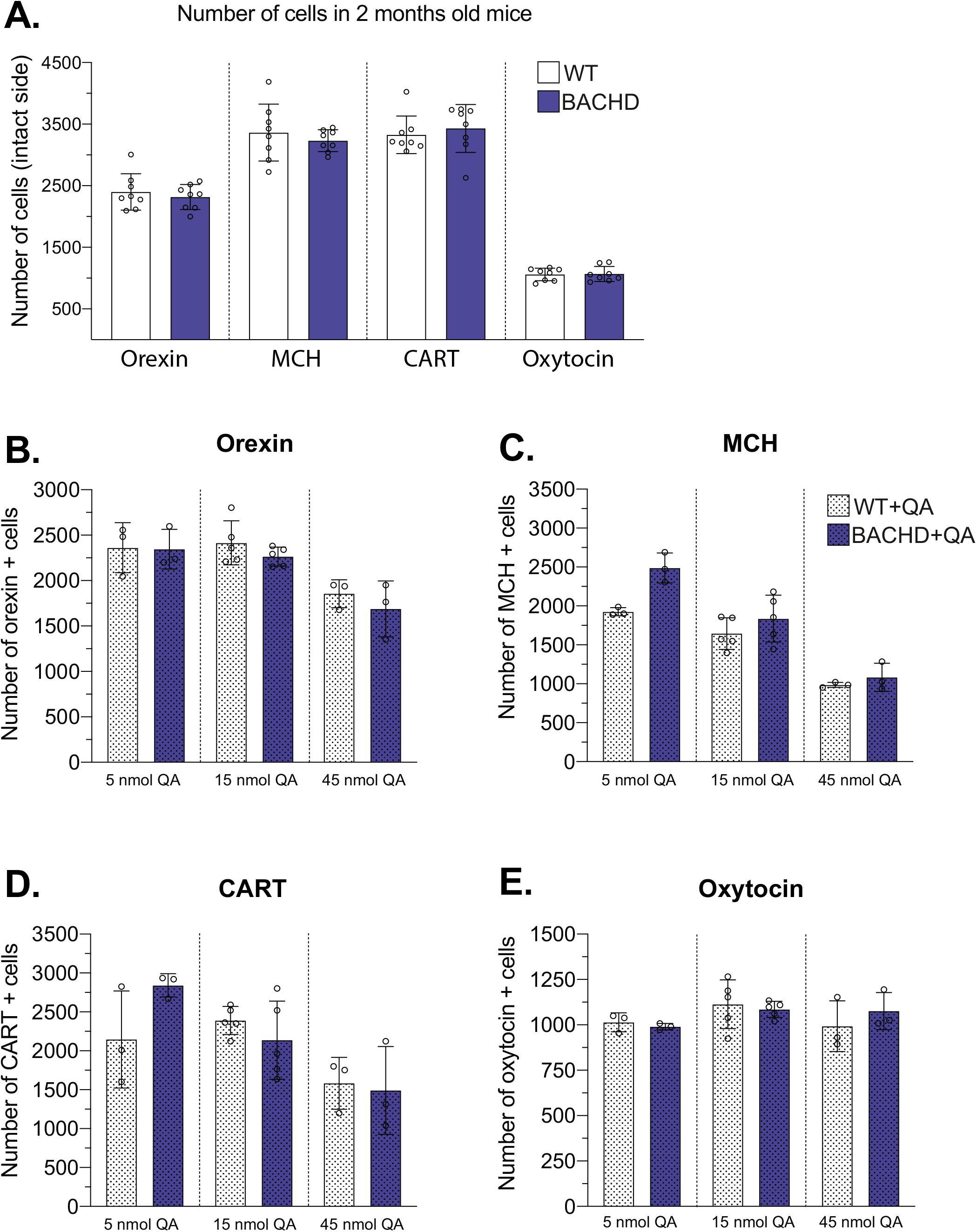
Assessment of neuronal populations in the BACHD mice after hypothalamic injections of QA. **(A)** The number of neurons expressing orexin, MCH, CART, and oxytocin in the hypothalamus was quantified in 2 months old BACHD and WT mice. BACHD and their WT littermates were unilaterally injected with 5, 15 or 45 nmol QA in the hypothalamus at two months of age, and the hypothalamic neuropeptide expressing cells were examined at one-week post-injection. There was no significant change in the number of **(B)** orexin, **(C)** MCH, **(D)** CART, and **(E)** oxytocin expressing cells in response to QA injection in BACHD mice compared to WT mice. Data are represented as scatter dot plots, and bars represent mean ± SEM.

## Conclusions

Excitotoxicity has been proposed to play a role in the selective vulnerability of MSNs in HD. Transgenic HD mouse models show a biphasic alteration in sensitivity to excitotoxicity in the striatum. There is also a selective loss of hypothalamic neurons in HD. However, exposure to excitotoxicity through injections of QA in the hypothalamus of mice does not fully recapitulate the hypothalamic neuropathology in clinical HD. The transgenic R6/2 and the BACHD mice do not display altered sensitivity to the excitotoxin QA as compared to their controls. Taken together, our data do not provide support for a role of excitotoxicity in the selective vulnerability of hypothalamic neurons in HD.

## Acknowledgements

This work was supported by grants from the Swedish Research Council to ÅP (2013/03537 and 2018/02559), the Province of Skåne State Grants (ALF) to ÅP and the Knut and Alice Wallenberg Foundation to ÅP (# 2019.0467). RSK was supported by Svenska Sällskapet för Medicinsk Forskning fellowship. We thank Björn Anzelius, Anneli Josefsson, Anna Hansen, Ulla Samuelsson and Ulrika Sparrhult-Björk at Lund University for their valuable technical assistance.

## Abbreviations

ANOVA: analysis of variance
BACHD: bacterial artificial chromosome mouse model of HD
CART: cocaine and amphetamine-regulated transcript
DAB: 3,3’-diaminobenzidine
HD: huntington disease
HTT: huntingtin
LHA: lateral hypothalamic area
MCH: melanin-concentrating hormone
MSN: medium spiny neuron
NMDA: N-methyl-D-aspartate
PFA: Paraformaldehyde
PVN: Paraventricular nucleus
QA: quinolinic acid
ROI: region of interest
RT: room temperature
SEM: standard error of mean
TBS: tris-buffered saline
TBS-T: TBS-triton-X
TH: tyrosine hydroxylase
WT: wild type
YAC: yeast artificial chromosome

## Notes

### Competing Interest Statement

The authors have declared no competing interest.

